# Multiplex analysis of cytokines and chemokines in persons aging with or without HIV

**DOI:** 10.1101/2023.01.30.526135

**Authors:** Kyle W. Kroll, Griffin Woolley, Karen Terry, Thomas A. Premeaux, Cecilia M. Shikuma, Michael J. Corley, Scott Bowler, Lishomwa C. Ndhlovu, R. Keith Reeves

## Abstract

People with HIV (PWH) on combined antiretroviral therapy (cART) are living longer lives due to modern cART advances and increased routine medical care. The full landscape of aging with HIV is unclear; given that HIV emerged relatively recently in human history and initially had a high mortality rate, there has not been a substantially aged population to evaluate. In the present study, we set out to perform high throughput plasma analyte profiling by multiplex analysis, focusing on various T helper (Th)-related cytokines, chemokines, and pro- and anti-inflammatory cytokines. The primary goals being to provide reference ranges of these analytes for aging PWH cohorts, as well as testing the utility of high throughput multiplex plasma assays. The cohort used in this study was comprised of age-matched healthy donors (aged 32.6-73.5), PWH on cART (aged 26.7-60.2), and viremic PWH (aged 27.5-59.4). The patients in each group were then stratified across the age span to examine age-related impacts of these plasma biomarkers. Our results largely indicate feasibility of plasma analyte monitoring by multiplex and demonstrate a high degree of person-to-person variability regardless of age and HIV status. Nonetheless, we find multiple associations with age, duration of known infection, and viral load, all of which appear to be driven by either prolonged HIV disease progression or long-term use of cART.

## Introduction

Combination antiretroviral therapy (cART) has dramatically improved the prognosis of people with HIV (PWH) due to effective viral suppression and CD4^+^ T cell recovery[1]. Increased survival rate, along with increased incidences of HIV acquisition among people over 50 years old, is now shifting the population of PWH to an older demographic[2]. While aging alone causes an accrual of alterations throughout the body over time, which are associated with a variety of chronic diseases and potential changing susceptibility to infectious diseases[3, 4], the compounded effect of long-term HIV infection and associated chronic inflammation in PWH as they age is not fully understood [5, 6]. Chronic immune dysfunction associated with HIV infection may exacerbate age-related comorbidities and contribute to early onset of aging-related chronic conditions[7-9] and emerging data support contribution that cART itself may influence the aging process[7]. Pre-exposure prophylaxis has been proven the most effective HIV prevention method for high-risk individuals such as the aging population, although this population is also at a higher risk of developing side effects from exposure to the active ingredient tenofovir[10]. With access to adequate treatment and care, the average life expectancy of PWH is now comparable to those without HIV infection, although management of comorbidities remains a complex issue[11-13]. Biological changes in aging are often reflected in the plasma proteome, where plasma sampled at different stages of life reveal a variance in biological pathways and associations with age-related diseases and phenotypic traits[4]. Indeed, plasma proteomics is an established source for early biomarkers of systemic disease[14], and has previously revealed higher inflammatory protein concentrations within PWH on long-term cART compared to healthy donors (HD), including mucosal defense chemokines[15]. Both viremic and virally suppressed PWH have shown age-related monocyte activation markers comparable to older HD[16]. To determine whether immune-related proteins are impacted by HIV across the age spectrum, we conducted a high-throughput multiplex immunoassay[17] to measure cytokines and chemokines in the plasma from HIV uninfected donors, PWH with viral suppression by cART, and viremic PWH over a wide age distribution.

## Results

### Cohort demographics

The cohort used for this study was comprised of three major age-mapped groups: HIV uninfected donors (HD; n = 16; aged 32.6-73.5), PWH on cART (cART; n = 20; aged 26.7-60.2), and PWH off cART and with viremia (Viremic; n = 14; aged 27.5-59.4). Viremic patients were off cART and had viremia at the time of sample acquisition. The median ages of each group were comparable (**Supplemental table 1**). Other demographic information such as percent male, CMV IgG titers, viral load measures, CD4 counts, and duration of known infection are available in **Supplemental Table 1**.

### Analyte concentrations

Most analyte concentration measurements were not significantly different between groups, but clear observable trends were informative to provide expected reference ranges for each of the analytes measured (**Fig. 1-4; Supplemental Table 2)**. For the purposes of data analysis, analytes were separated into groupings based on similarity of action *in vivo*[18]: CD4^+^ T helper (Th) lymphocyte types 1 and 2 (Th1/Th2) associated cytokines (**Fig. 1**), Th9/Th17/Th22 and regulatory T cell (Treg) associated cytokines (**Fig. 2**), inflammatory cytokines (**Fig. 3**), and chemokines (**Fig. 4**). Concentrations of Interleukin 2 (IL-2), IL-6, IL-8, tumor necrosis factor alpha (TNF-α) (**Fig. 1**), and IL-15 (**Fig. 3**) were consistently higher among individuals aged over 45 compared to those aged under 45 for all groups.

**Fig 1.**
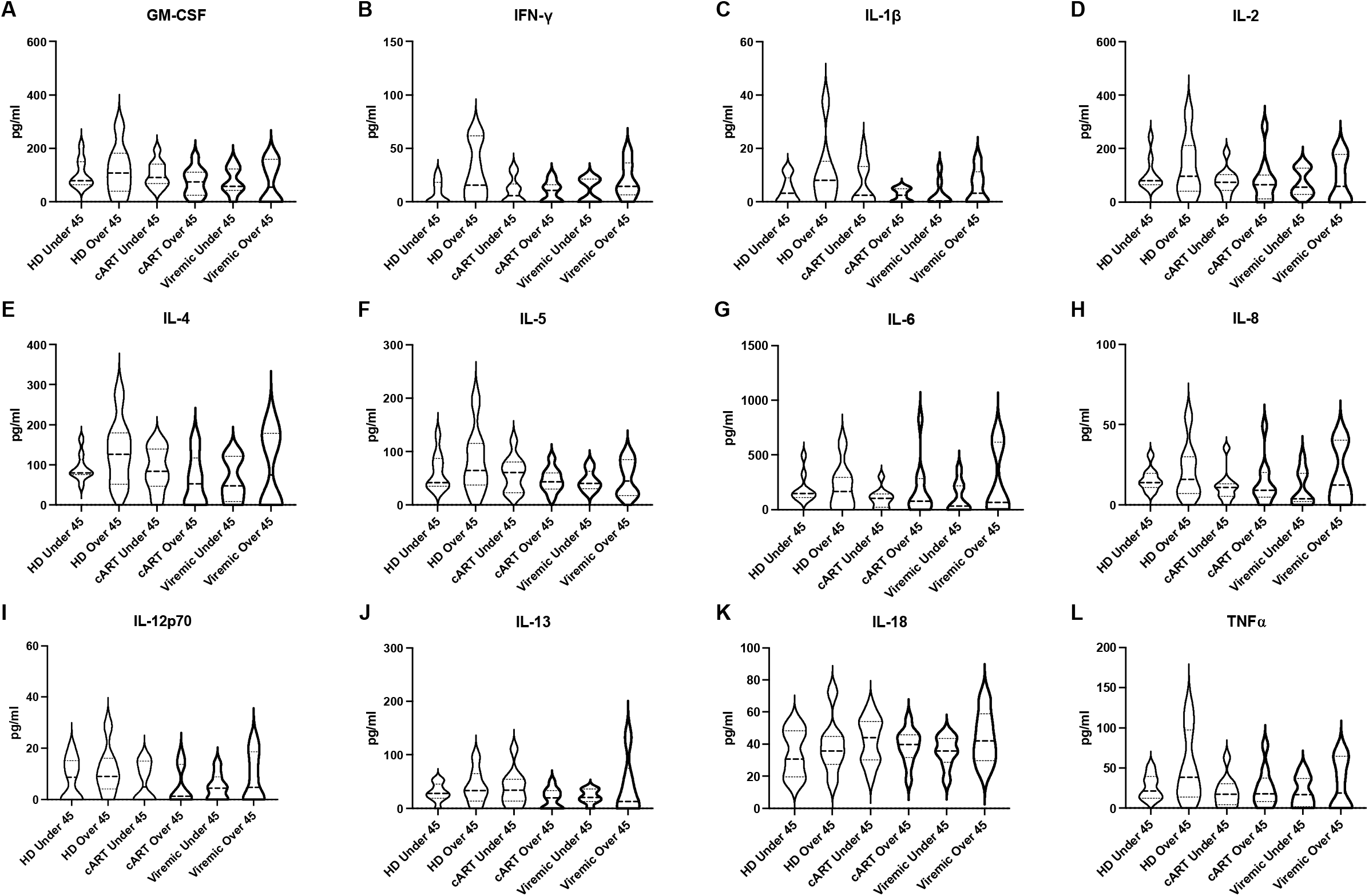

**Fig 2.**
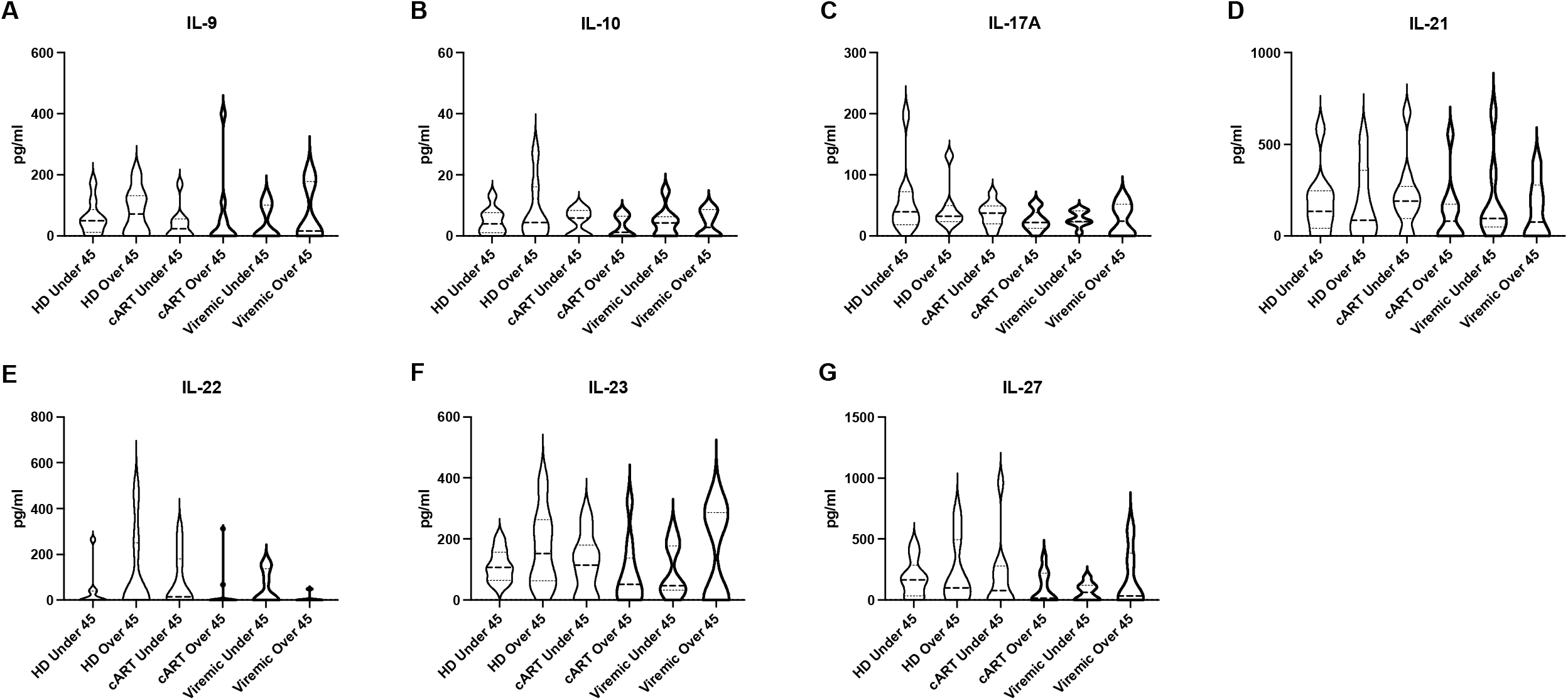

**Fig 3.**
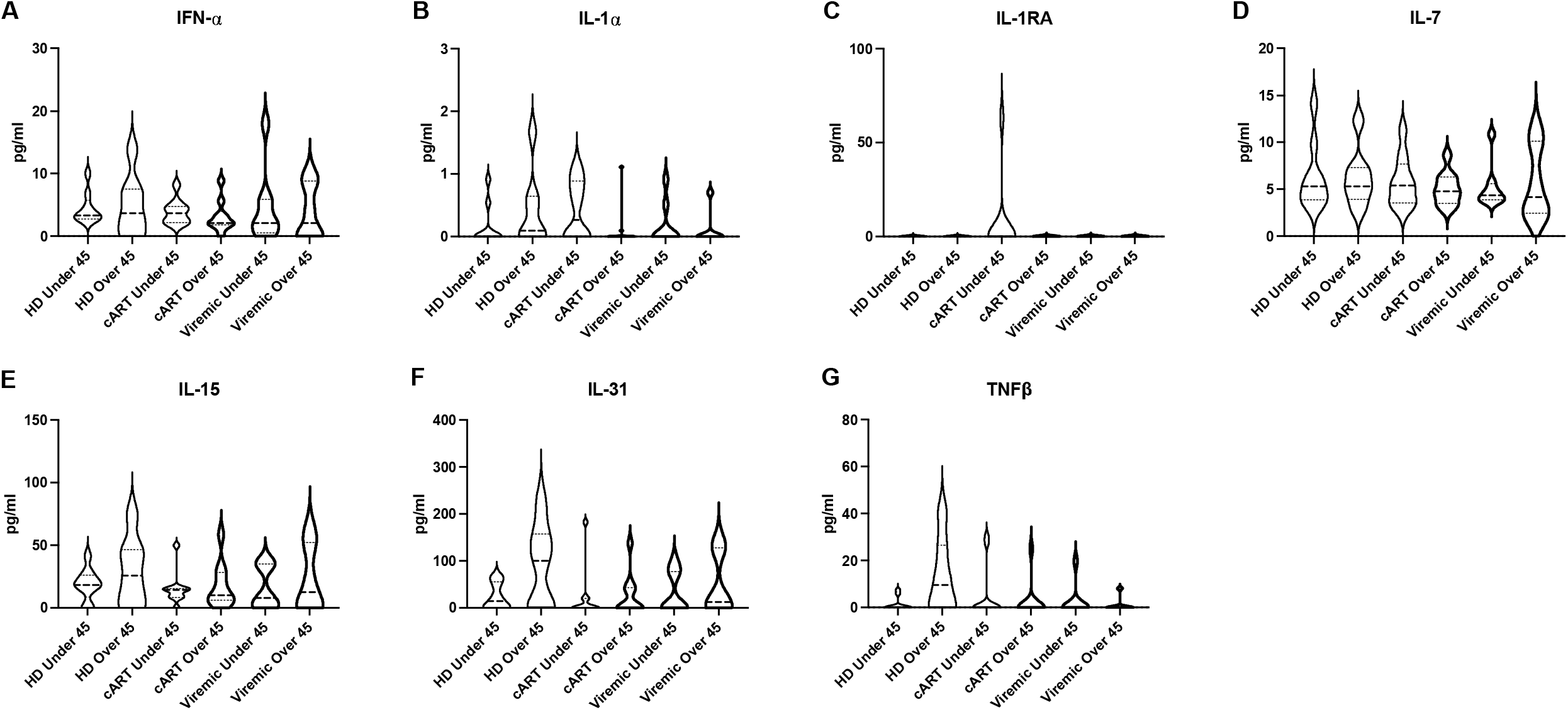

**Fig 4.**
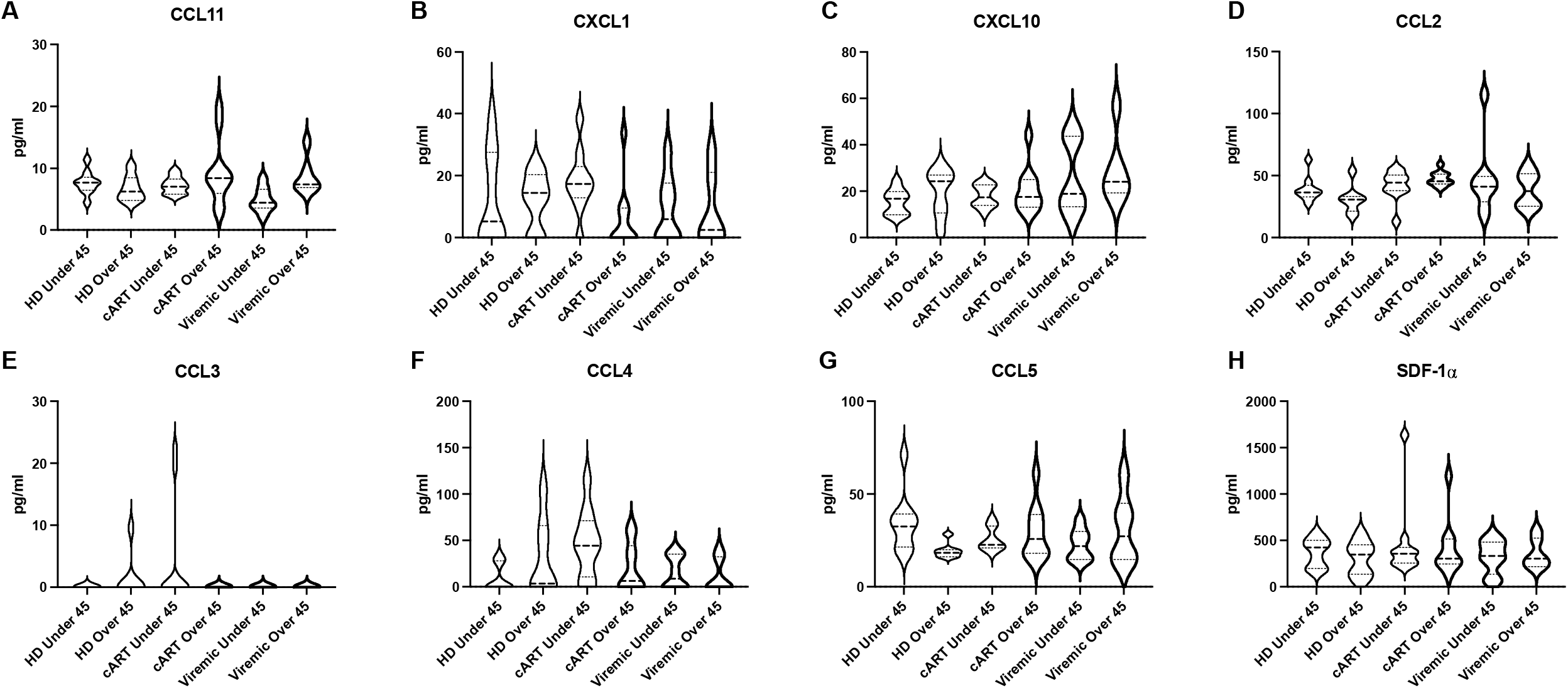

### Analyte correlations

HIV infection has been shown to have wide ranging system changes throughout infection[19-21]. To determine possible clinical correlations to the analytes in our study, we performed Pearson correlations between each analyte and CD4^+^ T cell counts, duration of known infection, and viral loads in viremic patients (**Table 1**). Of note, we observed several analytes which had significant correlations with duration of known infection in the cART group, including CCL11 (R = 0.53, *p* = 0.0166), granulocyte macrophage colony-stimulating factor (GM-CSF) (R = -0.46, *p* = 0.0403), IL-1*α* (R = -0.49, *p* = 0.0277), and IL-4 (R = -0.45, *p* = 0.0447). These findings are not entirely unexpected due to previous reports often showing pathologically lower levels of some cytokines in treated PWH compared to untreated PWH[22].

CCL11 also had a significant correlation with age among viremic patients (R = 0.62, *p* = 0.0176) (**Table 1**). Macrophage inflammatory protein-1 alpha (MIP-1α/CCL3) also had a significant correlation with age, in this case among PWH on cART (R = -0.55, *p* = 0.0111) (**Table 1**). Surprisingly, few correlations in viremic PWH are significant or show moderate trends when examining viral loads (VL). The chemokine C-X-C motif ligand 1 (GRO*α*/CXCL1) had a significant correlation with HIV VL (R = 0.69, *p* = 0.0063), and was also significantly correlated with age within patients on cART (R = -0.56, *p* = 0.0109) as well as with CD4^+^ T cell counts within viremic patients (R = -0.74, *p* = 0.0027) (**Table 1**). Otherwise, IL-2 (R = -0.46, *p* = 0.0434) and IL-8 (R = -0.47, *p* = 0.0385) each had significant correlations with CD4^+^ T cell counts among PWH on cART (**Table 1**). Taken together, our findings suggest that analyte correlations in PWH on cART were modulated differently compared to viremic PWH.

### Analyte pathway analysis

Heatmap visualization of analyte concentrations (**Fig. 5**) highlights the variance by samples for all analytes. While the degree of variability was high, as shown above in correlative analyses some associations do appear for analytes with higher expression in individuals over 45 group for all groups. To get a better understanding of underlying expression patterns, analyte pathway analysis was subsequently performed (**Fig S3**). In PWH on cART over 45, there is an upregulation compared to age-matched HD in the “Viral protein interaction with cytokine and cytokine receptor”, and more specifically upregulations in the chemokines MCP-1/CCL2, CCL11, and RANTES/CCL5. These indicate a potential increase in immune trafficking, but whether these trafficking are to sites of HIV reservoir or other opportunistic infections related to age is currently unknown. In addition, several Th1/Th2 related cytokines: Interferon-gamma (IFN-γ), IL-4, IL-5, IL-12A, and IL-13 are all downregulated compared to HD. Similar analysis was performed for viremic PWH, although we did not observe similar results (data not shown), which may be a result of lower statistical power due to sample distributions. These findings suggest that in PWH on cART exhibit increased immune trafficking but lower inflammation, consistent with previous studies[23, 24].

**Fig 5.**
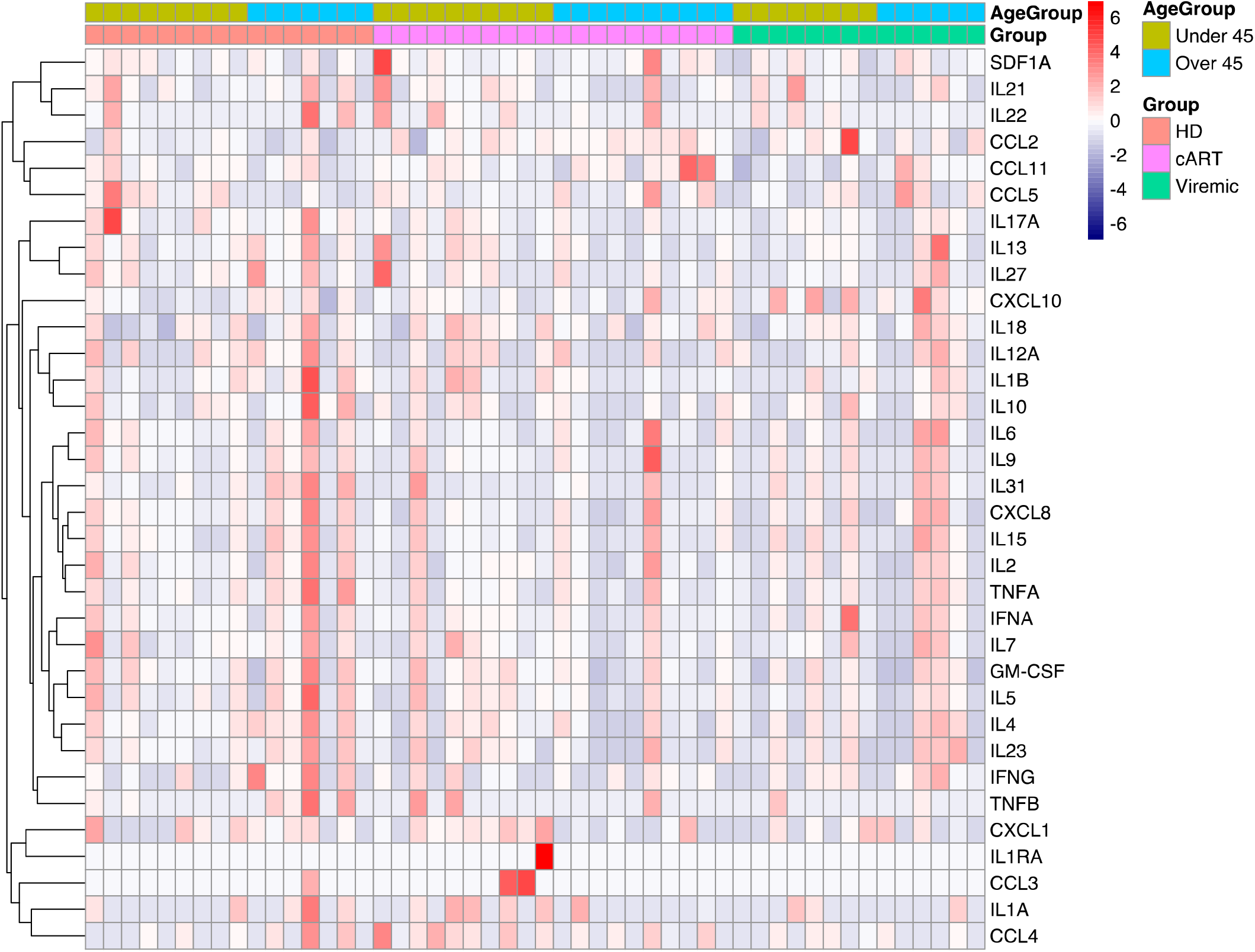

Principal Component Analysis (PCA) was performed using analyte concentrations without any input labels for samples (sample ID, groups, and age). Label-free classification allows the samples to cluster based entirely on expression patterns instead of using classification labels as guides. After PCA reduction, group and age group labels were attached (**Fig. 6**). PCA analysis reveals that across age groups and HIV status, samples do not clearly cluster by any combination of age and HIV status (**Fig. 6**). However, there are two major clusters indicating potential for two distinct analyte profiles among subjects. PCA loadings are comprised of eigenvectors (direction) and eigenvalues (magnitude) of influence from each of the input variables. The direction and magnitude for each variable can be plotted to see their relative impact on the output PC dimensions. Upon examination of PCA loadings, this separation is driven more by analyte expression, high vs low, in the samples and that the cluster with higher expression appears to favor samples from the cART group. These results are consistent with prior studies[24] which show that even though cART use reduces the chronic inflammation of HIV, it does not return it to levels seen in healthy controls. Notably, the analytes

**Fig 6.**
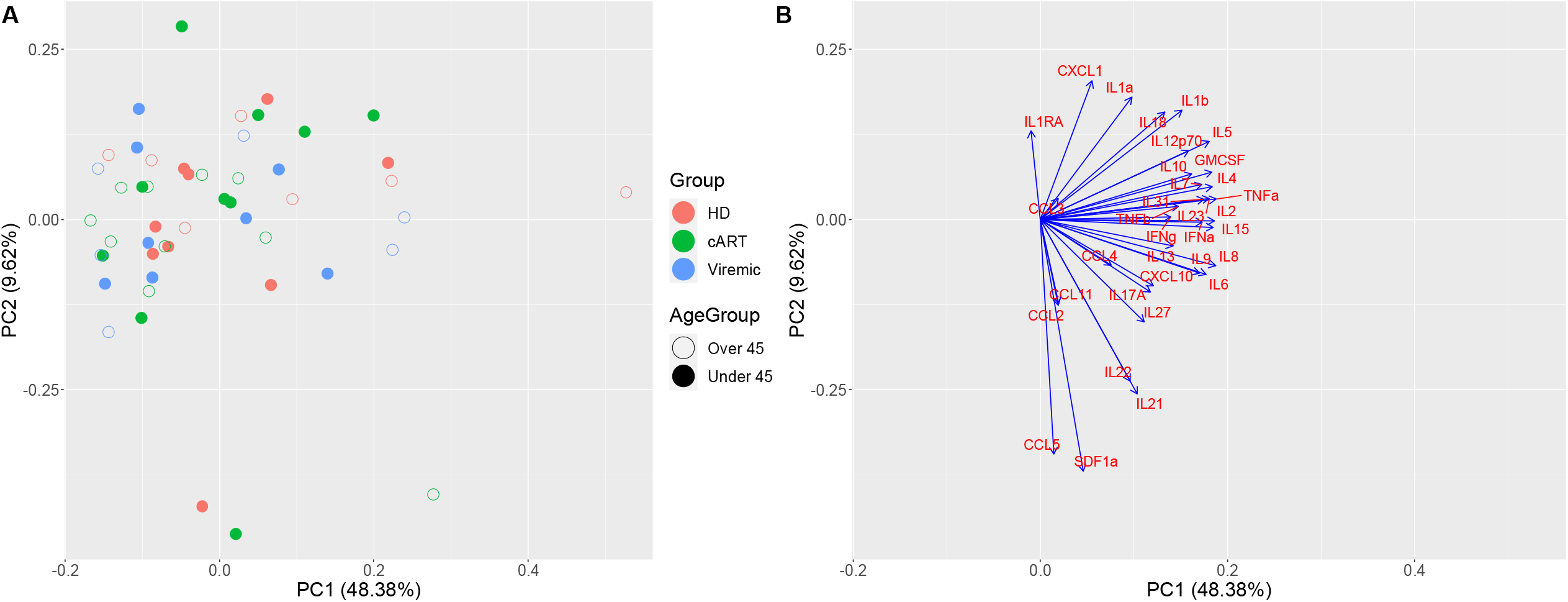

IL-2, TNF-α, IL-15, and several other interleukins have major impacts on the first principal component. While IL-1RA, RANTES/CCL5, and SDF-α are major driving factors for the second principal component. Based on these loadings, the first PC is largely driven by interleukin expression, while the second PC is largely driven by chemokine expression, highlighting the fact that inflammatory cytokine expression is still a major driver in PWH on cART separation from the other groups, regardless of age.

## Discussion

Parallels between natural aging and the premature aging that corresponds with HIV have been established[25], and biomarkers with inflammation and immune activation may be associated with the accelerated aging observed during HIV infection[25]. Additionally, HIV elite controllers have been observed to have higher levels of soluble inflammatory markers compared to HIV-suppressed or uninfected individuals[26], and elevated levels of soluble biomarkers has been associated with occurrences of non-AIDS-defining events in PWH on cART[27]. There are many different types of cytokine profiles, each with critical roles in the immune response[28]. In this study we examined profiles encapsulating Th1/Th2 helper T cells, Th9/Th17/Th22/Treg helper T cells, inflammatory cytokines, and chemokines. Th1/Th2 helper T cells have polarizing cytokine production profiles with an array of different immune responses[29]. The Th9/Th17/Th22/Treg class of helper T cells have activity which has been associated with immune balance and function in the context of HIV infection and its progression to AIDS[30]. Inflammatory cytokines have long been recognized for their vital role in protection against pathogens, although inappropriate activation can lead to acute and chronic disease[18, 31], and chronic inflammation has been associated with increased risk of disease in aging people[32]. Chemokines play vital roles as regulatory molecules in the function and trafficking of leukocytes, lymphocytes, and lymphoid tissues[33-35]. HIV infection manipulates these functions to facilitate its own transmission and pathogenesis[36]. In combination, cytokines and chemokines regulate a healthy immune response throughout the body when functioning properly[37, 38]. However, the dysregulation of cytokines and chemokines which occurs during HIV infection can increase pathogenesis by contributing to the depletion of CD4^+^ T cells and the elevation of VL[39]. Our focus in this study is to examine analyte expression profiles across three groups: HD, PWH on cART, and PWH with viremia. Subsequently we examined the impact of age for each of these groups by examining expression profiles comparing those under 45 and over 45 years of age. In addition, we also examined correlation of analyte expression with several clinical data points: age, CD4^+^ T cell count, duration of known infection, and viral loads.

Of the Th1/Th2 analytes, IL-2 and IL-8 were both observed to have a significant negative correlation with CD4^+^ T cell counts among PWH on cART. IL-2 is a secretory cytokine that can stimulate T cells, B cells, and NK cells, and has in its recombinant form been utilized as a therapeutic for PWH concurrent with cART due to its ability to slow the apoptosis of HIV-infected cells[40, 41]. Our findings with IL-2 indicate that PWH on cART may experience an increase in immune activation and proliferation, or the compensatory function of IL-2 as CD4^+^ T cell counts decrease. IL-8 is a proinflammatory cytokine which has been identified as a marker of chronic inflammation within PHW on cART[42], and blocking IL-8 in tandem with GRO*α*/CXCL1 has been demonstrated to inhibit HIV replication within T cells and macrophages[43]. Additionally, IL-8 and IFN-γ and each trended negatively with duration of known infection among viremic individuals. IFN-γ is a cytokine secreted by activated T cells and NK cells with a wide range of immune response functions, such as macrophage activation, antiviral and antibacterial immunity mediation, and several other important immunoregulatory functions[44]. IFN-γ has been observed in higher concentrations within PWH compared to HD[45], and PWH with high levels of circulating IFN-γ exhibited a lower CD4^+^ T cell count recovery after 4 years of being on cART[46]. Additionally, IFN-γ trended positively with age among HD. This supports findings by past researchers that the production of IFN-γ is directly correlated with age[32, 47].

The Th1/Th2 analytes GM-CSF and IL-4 each had a significant negative correlation with duration of known infection among PWH on cART. GM-CSF promotes myeloid cell development and maturation, dendritic cell differentiation and survival *in vitro*, and has a major role in the regulation of intestinal immune and inflammatory responses[48]. GM-CSF signaling in alveolar macrophages has been shown to be disrupted in HIV infection[49], which may explain our observation of its decline over longer durations of infection. An imbalance in GM-CSF production or signaling may contribute to harmful inflammatory conditions, potentially giving it a role in the concept of inflammaging and/or chronic inflammation seen in long term HIV infection. Meanwhile the cytokine IL-4 has been documented to regulate B and T cell growth, immunoglobulin secretion, eosinophile recruitment, and macrophage activation[50, 51]. We observed a negative correlation with both duration of known infection and CD4 counts in PWH on cART, which supports a previous finding where serum levels of IL-4 were significantly lower in PHW on sustained HAART compared to before commencement of treatment[52]. Furthermore, IL-12p70 was found to be negatively trended with duration of known infection among PWH on cART. IL-12p70 is the bioactive form of IL-12, comprised of p35 and p40, and plays a role in the induction of IFN-γ production[53, 54]. IL-12p70 is only transiently increased in viremic individuals compared with HD in correlation with VL, and its response to HIV infection may be overshadowed by other cytokines such as IL-23[54].

Among the Th9/Th17/Th22/Treg analytes, we observed several interesting trends. IL-9 trended negatively with duration of known infection among viremic individuals and IL-10 trended negatively with CD4^+^ T cell counts among PWH on cART. Like IL-4, IL-10 is a proinflammatory cytokine that facilitates the differentiation of naïve CD4^+^ T Cells into Th2 subtypes[55]. IL-10 has been observed to significantly and gradually decrease within PWH on highly active antiretroviral therapy (HAART). IL-22 was found to be negatively trended with age among viremic individuals. This can be explained by the fact that HIV infection depletes mucosal Th22, Th17, and innate lymphoid cells, which are responsible for the production of inflammatory cytokines such as IL-22, although this deficiency can be rapidly restored by the administration of cART[56]. IL-23 was found to be negatively trended with duration of known infection among PWH on cART. The inhibition of IL-23 signaling and expression have been reported within PHW on long-term HAART[57, 58].

Regarding inflammatory cytokines, IL-1α had a significant negative correlation with duration of known infection among PWH on cART. IL-1α induces proinflammatory effects and is central to the pathogenesis of numerous conditions characterized by body-wide organ or tissue inflammation[59]. Our findings here could be partially explained by the ability of cART to reduce systemic inflammation during acute HIV infection[60]. IL-31 trended negatively with duration of known infection among viremic individuals. IL-31 is involved in maintaining hematopoietic stem cells and influencing proinflammatory activities of macrophages, monocytes, and T cells[61, 62]. Serum concentrations of IL-31 have been reported to be higher in viremic patients compared to HD[62]. However, as IL-31 is primarily produced by CD4^+^ T cells[63, 64], and CD4^+^ T cells are depleted during HIV infection[65, 66], additional research is necessary to fully understand the potential relationship between IL-31 function and HIV infection. TNF-β was found to be positively trended with viral load among viremic individuals. TNF-β has been reported to stimulate HIV replication within T cells and monocyte-derived macrophages[67].

The last analyte grouping we analyzed were chemokines, among which CCL11 was found to be significantly positively correlated with age among viremic individuals, although it trended negatively with age among HD, indicating that it is impacted by some factor or combination of factors resulting from HIV infection. CCL11 plays a role in a skewed immune response toward a Th2 profile[68]; Th2-type cytokines are associated with anti-inflammatory responses to mediate the proinflammatory responses of Th1-type cytokines[69]. In addition to its role in the immune response, CCL11 has been demonstrated to have associations with aging, neurogenesis and neurodegeneration, and microglia[70]. CCL11 also has a significant positive correlation with duration of known infection among PWH on cART. High plasma levels of CCL11 have been related to a decrease in CD4^+^ T cells in HIV elite controllers with sustained viral suppression[71]. GRO*α*/CXCL1 was found to be significantly negatively correlated with both age among PWH on cART, and with CD4^+^ T cell counts among viremic individuals. GRO*α*/CXCL1 has a crucial role in inflammation[72], and has been shown to have potential as a marker of disease activity[73]. GRO*α*/CXCL1 is among a family of IL-8-related chemokines which have been observed to be upregulated when exposed to HIV-infected T cells[74]. Perhaps the upregulation of GRO*α*/CXCL1 initiated by HIV-infected T cells continues even as the T cells are depleted by HIV disease progression[66]. GRO*α*/CXCL1 was also observed to have a significant positive correlation with viral load among viremic individuals, further demonstrating its ability to be used as an indicator of disease activity[73]. Meanwhile, IP-0/CXCL10 and MCP-1/CCL2 were both found to be positively trended with viral load among viremic individuals. Like GRO*α*/CXCL1, IP-10/CXCL10 has also been identified as a robust soluble biomarker of monocyte activation with potential as a prognostic marker for HIV, as levels have been observed to increase along with VL[75, 76]. MCP-1/CCL2 is a proinflammatory chemokine which supports the replication of HIV[77] and has been observed within aging PWH at significantly higher levels in comparison to age-stratified HD[78].

MIP-1α/CCL3 was found to be significantly negatively correlated with age among PWH on cART. MIP-1α/CCL3 is a macrophage-derived chemotactic chemokine with proinflammatory activities and a role in the effector immune response which has been recognized as a potential marker for the detection of various diseases[79, 80]. IP-10/CXCL10 trended negatively with CD4^+^ T cell counts among PWH on cART. IP-10/CXCL10 has been identified by other researchers at higher levels within PWH who have confirmed lowered CD4^+^ T cell counts[76]. MIP-1β/CCL4 was found to be positively trended with duration of known infection among viremic individuals. MIP-1β/CCL4 is proinflammatory cytokine which has been observed to have a significant positive correlation with plasma levels of the antiviral drug emtricitabine among individuals taking PrEP[81]. RANTES/CCL5 was found to be positively trended with duration of known infection among PWH on cART. RANTES/CCL5 is expressed by inflammatory cells such as T cells and monocytes, and its interaction with the CCR5 receptor has been targeted as a strategy for long-term control of HIV infection[82, 83]. These findings provide supportive evidence that a longer duration of known infection, and, indirectly, a longer duration of cART administration, corresponds with a reduction in inflammation markers.

Of all the analytes measured, concentrations of IL-2, IL-6, IL-8, IL-15, and TNF-α were higher among individuals aged over 45 compared to those younger than 45 across groups. Raised levels of IL-6 have been associated with age-related disease[84-93], and IL-6 has also been considered as a potential as a biomarker of clinical outcomes from HIV-related inflammation[94]. Additionally, PWH receiving treatment have higher IL-6 levels than those who do not, and elevated IL-6 levels in PWH on cART are associated with advanced age and elevated body mass index[90, 94]. The over-activation of TNF-α signaling has been associated with chronic inflammation and the eventual development of pathological complications[95]. TNF-α has been previously recorded within PWH taking long-term suppressive cART at levels comparable to those of elderly PWH who did not survive infection, and this may be due to persistent alteration of TNF-α levels in PWH causing tissue damage and influencing the development non-AIDS-defining illnesses while the replication of HIV itself is being controlled by cART[96]. IL-15 has established roles in the development and maintenance of NK and CD8^+^ T cells and has been observed to be significantly elevated within viremic individuals compared to HD[97, 98]. Perhaps our observed increase in IL-15 could be the result of a natural attempt to overcome the reduced functionality of NK cells in aging adults[99].

Overall, we observed several significant correlations as well as nonsignificant trends between analytes and the conditions of age, CD4^+^ T cell counts, duration of known infection, and viral load. As this data was acquired from plasma, it is more representative of systemic events, but can still be indicative that trafficking modulation plays a large role during HIV infection. Based on our findings, there may be moderate impacts on the Th1/Th2 cells, specifically inflammation from GM-CSF and cell survival from IL-2 and IL-4. While the sample size for this study was moderate, it may be underpowered to tease out subtle differences that may be masked by individual variability. However, these data serve as an important reference for future studies seeking to examine changes in the immune landscape during aging with or without HIV. Our data further enlightens on the dynamics of the aging process with HIV, particularly immunological changes, that could help guide future strategies to address the complex clinical care that comes with this shifting age demographics of the PWH population.

## Methods

### Study Participant Characteristics

Cryopreserved human peripheral blood mononuclear cells (PBMCs) were obtained from the Hawai’i Aging with HIV-1 Cohort (HAHC) at University of Hawai’i[102]. Details of the HAHC study enrollment and clinical characterization were previously published[103] and approved by the University of Hawai’i Institutional Review Board #10675. All participants signed institutional review board–approved informed consent forms prior to participation. CD4^+^ T lymphocyte counts were obtained in real-time by standard technique from a local CAP certified reference laboratory. Demographic and clinical details of study participants are provided in **Table S1**. Samples were grouped into three groups: healthy donors (HD; n=16), PWH on cART and virally suppressed (cART; n=20), and PWH off cART with viremia (Viremic; n=20).

### CMV antibody titer quantification

Matched cryopreserved plasma samples were used for quantification of Human Cytomegalovirus (CMV) IgG and IgM antibody titers (Quest Diagnostics; order code 6732; Marlborough, MA, USA) as previously published[102].

### Multiplex assay determination of plasma cytokine/chemokine concentrations

As a microsphere bead capture assay, the ProcartaPlex multiplex assays require only 25µL of plasma and takes only four hours to yield analyzed results[18, 104]. With Luminex multi-analyte profiling (xMAP) technology simultaneous analysis of up to 50 analytes per 20-50μL sample can be accurately performed[105], while an enzyme-linked immunosorbent assay (ELISA) generally requires a minimum sample volume of 100μL[106, 107]. As a precise method of plasma cytokine profiling, Luminex is more cost- and time-effective than ELISA and other immunoassays, making it ideally suitable for high-throughput biomarker identification[108, 109], especially in situations where sample availability is restricted.

Cryopreserved plasma samples underwent no more than 2 freeze-thaw cycles prior to analysis. For this assay, plasma samples were taken from -80°C storage and thawed at room temperature (RT) before being deactivated for viral contaminates by diluting 50/50 with 2% triton for 1 hour at RT. For generation of standard curves, manufacturer-provided lyophilized standards were reconstituted and prepared by 4-fold serial dilutions for a total of 8 standards as described in the kit insert. These standards were analyzed in duplicate on a 96-well optical plate alongside the prepared plasma samples which were also ran in duplicate. Two additional wells were run without standards or samples to provide background measurements. The concentrations of plasma cytokines and chemokines were measured via Luminex xMAP technology utilizing the Procartaplex Human Cytokine/Chemokine Convenience Panel 1A 34-plex (Thermo Fisher EPXR340-12167-901) in accordance with the manufacturer’s instructions.

Briefly, 50μL of vortexed magnetic capture beads were added to each well and placed on a handheld magnet to collect beads at the bottom of the well. The plate was then washed with provided wash buffer (WB) twice. 50μL of samples were then added in duplicate to assigned wells. For standards, 25μL of each standard plus 25μL of Universal Assay Buffer (UAB) was added to the appropriate wells. Duplicate background wells had 50μL of UAB added. The plate was then sealed, placed on a shaker, and incubated at 4°C for 20 hours. The plate was next washed twice, with WB and 25μL of detection antibody added to each well, and shaken for 30 minutes at RT. The plate was then washed twice and 50μL of Streptavidin-PE was added to each well and incubated at RT on the shaker for 30 minutes. Finally, the plate was washed twice, 120μL of Reading Buffer added to each well, and then the plate was incubated on shaker at RT for 5 minutes. Analysis of the plate wells was performed using the Luminex 200 (Luminex Corporation) which was calibrated, and performance was validated as per instrument validation protocol. Measurements were reported with the xPONENT 4.2 software (Luminex Corporation).

### Bioinformatics analyses

Final concentrations of analytes were loaded into R v4.1.2[110]. Fold change values were calculated by first taking the mean expression of each analyte for each group and age group. Mean expression values were then divided by the matched age group of the HD cohort and the resulting fold changes were transformed with the log2 function. *P*-values were determined using two-tailed t-tests. Term enrichment analysis was performed with the pathfindR package[111].

### Statistical tests

Statistical tests and violin plots were performed in GraphPad Prism v9.4.0. *P*-values were calculated using ordinary one-way ANOVA followed by adjustment with Tukey’s multiple comparison test. Spearman correlation was performed in GraphPad Prism for correlation analyses between analytes and clinical markers such as age, duration of known infection, CD4^+^ T cell counts, and viral loads.

## Data availability

The data generated for this study is available upon request to the corresponding author.

## Author contributions

R.K.R. and L.C.N. designed the study. M.J.C., T.A.P., S.B., and C.M.S. facilitated sample evaluation and metadata. K.T. carried out Luminex assays. G.W. and K.W.K. analyzed data and prepared the manuscript.

