## Supplemental Materials for "Multiplex analysis of cytokines and chemokines in persons aging with or without HIV"

|  | **Median age (range)** | **% Male** | **Median CMV IgG (range)**  **(U/mL)** | **Median log10 VL (range) (RNA copies/mL)** | **CD4 Count (range)**  **(count/mL)** | **Duration of Known Infection**  **(years)**  **(range)** |
| --- | --- | --- | --- | --- | --- | --- |
| **HD**  **n=16** | 42.4  (32.6 - 73.5) | 81.25% | 5.25  (0.6 - 10) |  |  |  |
| **cART**  **n=20** | 48.2  (26.7 - 60.2) | 75.00% | 10  (5.8 - 10) |  | 502.5  (132 – 1380) | 11.1  (1.7 – 24.67) |
| **Viremic**  **n=14** | 42.3  (27.5 - 59.4) | 60.00% | 10  (0.6 - 10) | 4.39  (4.11 - 5.39) | 429.5  (10 – 749) | 5.39  (0.25 – 22.38) |

**Table S1. Cohort Demographics and Clinical Information.** Demographic and clinical information for cohorts of healthy donors (HD), people with HIV (PWH) on combination antiretroviral therapy (cART), and viremic individuals. Information includes median age, percentage male, median cytomegalovirus (CMV) immunoglobulin G (IgG), median log10 viral load (VL), CD4 counts, and duration of known infection.

|  | **HD** | | **cART** | | **Viremic** | |
| --- | --- | --- | --- | --- | --- | --- |
|  | ***Under 45*** | ***Over 45*** | ***Under 45*** | ***Over 45*** | ***Under 45*** | ***Over 45*** |
| Eotaxin (CCL11) | 7.67 (4.49 - 11.38) | 6.2 (4.66 - 10.32) | 7.04 (5.69 - 9.7) | 8.45 (3.07 - 19.98) | 4.42 (2.14 - 8.36) | 7.41 (6.44 - 14.25) |
| GM-CSF | 79.23 (63.1 - 209.44) | 107.43 (0 - 282.06) | 91.025 (42.6 - 194.32) | 75.335 (0 - 168.67) | 57.58 (0 - 150.94) | 54.925 (0 - 163.63) |
| GRO⍺ (CXCL1) | 5.27 (0 - 38.52) | 14.4 (0 - 23.22) | 17.35 (0 - 38.35) | 0 (0 - 33.76) | 5.86 (0 - 28.1) | 2.48 (0 - 27.91) |
| IFN⍺ | 3.29 (2.57 - 9.99) | 3.67 (0 - 13.8) | 3.69 (1.78 - 8.18) | 2.105 (0 - 8.86) | 2.105 (0 - 17.98) | 2.11 (0 - 9.7) |
| IFNγ | 0 (0 - 26.9) | 15.85 (0 - 62.62) | 5.98 (0 - 31.29) | 10.55 (0 - 25.16) | 9.845 (0 - 22.46) | 14.7 (0 - 46.74) |
| IL-1⍺ | 0 (0 - 0.91) | 0.09 (0 - 1.67) | 0.265 (0 - 1.12) | 0 (0 - 1.11) | 0 (0 - 0.91) | 0 (0 - 0.7) |
| IL-1β | 3.3 (0 - 12.11) | 8.01 (0 - 37.53) | 2.52 (0 - 20.61) | 2.485 (0 - 5.73) | 0.37 (0 - 12.79) | 3.17 (0 - 15.73) |
| IL-1RA | 0 (0 - 0) | 0 (0 - 0) | 0 (0 - 63.23) | 0 (0 - 0) | 0 (0 - 0) | 0 (0 - 0) |
| IL-2 | 79.62 (63.97 - 242.95) | 96.82 (5.54 - 331.37) | 73.46 (0 - 185.47) | 65.485 (0 - 282.97) | 54.51 (13.55 - 143.32) | 58.71 (0 - 191.09) |
| IL-4 | 79.79 (49.04 - 163.32) | 126.33 (33.87 - 274.81) | 84.09 (0 - 156.66) | 52.655 (0 - 169.19) | 47.23 (0 - 129.82) | 75.185 (0 - 206.93) |
| IL-5 | 42.22 (32.88 - 131.79) | 64.53 (13.42 - 194.97) | 61.245 (19.34 - 120.38) | 43.605 (12.37 - 87.93) | 40.255 (22.35 - 78.82) | 45.085 (17.5 - 95.4) |
| IL-6 | 145.02 (87.53 - 495.53) | 164.88 (0 - 601.94) | 100.41 (0 - 301.63) | 78.6 (0 - 826.52) | 31.505 (0 - 367.03) | 65.2 (0 - 664.66) |
| IL-7 | 5.29 (3.06 - 14.12) | 5.29 (2.7 - 12.31) | 5.4 (2.75 - 11.29) | 4.765 (2.8 - 8.61) | 4.345 (3.67 - 10.86) | 4.145 (2.37 - 11.29) |
| IL-8 (CXCL8) | 13.89 (7.38 - 30.86) | 15.83 (4.88 - 54.59) | 10.71 (0 - 35.42) | 9.15 (0 - 49.08) | 3.65 (0 - 29.26) | 12.345 (0 - 40.8) |
| IL-9 | 49.96 (0 - 172.73) | 71.63 (0 - 199.8) | 23.44 (0 - 168.95) | 0 (0 - 398.62) | 0 (0 - 127.67) | 15.23 (0 - 203.29) |
| IL-10 | 3.95 (0 - 13.03) | 4.36 (0 - 27.19) | 5.79 (0 - 9.65) | 1.205 (0 - 7.73) | 4.28 (0 - 14.59) | 2.805 (0 - 9.22) |
| IL-12p70 | 8.76 (0 - 20.24) | 9.05 (0 - 28.65) | 5 (0 - 16.48) | 1.28 (0 - 17.78) | 4.435 (0 - 14.1) | 4.8 (0 - 22.79) |
| IL-13 | 28.2 (0 - 53.8) | 32.78 (0 - 96.12) | 34.145 (0 - 110.94) | 19.595 (0 - 50.23) | 20.68 (0 - 37.77) | 13.235 (0 - 132.65) |
| IL-15 | 18.18 (0 - 41.33) | 25.57 (0 - 73.99) | 14.015 (0 - 49.75) | 9.72 (0 - 58.4) | 7.83 (0 - 35.43) | 12.37 (0 - 62.02) |
| IL-17A (CTLA-8) | 39.56 (13.5 - 197.36) | 32.35 (18.97 - 130.75) | 36.87 (2.4 - 69.49) | 21.565 (0 - 51.96) | 23.58 (5.41 - 42.51) | 24.215 (0 - 62.56) |
| IL-18 | 30.64 (15.93 - 53.31) | 35.87 (16.6 - 72.41) | 44.01 (19.68 - 62.96) | 39.715 (17.46 - 55.61) | 35.84 (17.46 - 46.65) | 41.935 (23.85 - 68.55) |
| IL-21 | 133.22 (0 - 583.11) | 85.28 (0 - 526.26) | 189.93 (0 - 674.65) | 79.77 (0 - 554.56) | 95.14 (0 - 670.89) | 76.045 (0 - 384.02) |
| IL-22 | 0 (0 - 264.69) | 0 (0 - 464.83) | 15.32 (0 - 309.02) | 0 (0 - 312.14) | 0 (0 - 155.02) | 0 (0 - 48.69) |
| IL-23 | 106.77 (44.52 - 203.89) | 152.71 (29 - 382.05) | 115.495 (0 - 282.4) | 52.24 (0 - 322.65) | 47.12 (0 - 226.16) | 130.88 (0 - 320.33) |
| IL-27 | 168.63 (0 - 451.39) | 101.81 (0 - 702.22) | 77.73 (0 - 962.58) | 17.885 (0 - 344.18) | 65.37 (0 - 188.11) | 34.34 (0 - 573.38) |
| IL-31 | 14.49 (0 - 64.61) | 99.95 (0 - 224.14) | 0 (0 - 182.1) | 0 (0 - 137.55) | 0 (0 - 99.28) | 11.755 (0 - 138.64) |
| IP-10 (CXCL10) | 16.81 (8.98 - 24.32) | 24.26 (2.76 - 30.04) | 17.295 (10.56 - 24.56) | 17.435 (10.35 - 44.09) | 18.76 (8.16 - 44.69) | 24.04 (15.83 - 56.72) |
| MCP-1 (CCL2) | 36.46 (27.14 - 62.8) | 30.75 (17.38 - 53.48) | 44.19 (12.82 - 55.15) | 45.58 (39.74 - 59.08) | 41 (17.01 - 115.27) | 37.505 (22.87 - 57.19) |
| MIP-1⍺ (CCL3) | 0 (0 - 0) | 0 (0 - 9.54) | 0 (0 - 22.38) | 0 (0 - 0) | 0 (0 - 0) | 0 (0 - 0) |
| MIP-1β (CCL4) | 0 (0 - 33.07) | 3.72 (0 - 105.57) | 44.4 (0 - 115.84) | 6.455 (0 - 61.57) | 8.8 (0 - 37.52) | 0 (0 - 39.18) |
| RANTES (CCL5) | 32.66 (17.23 - 71.18) | 18.43 (15.97 - 28.35) | 22.83 (18.38 - 36.95) | 25.91 (14.51 - 61.05) | 21.985 (13.67 - 36.54) | 27.255 (14.54 - 60.37) |
| SDF-1⍺ | 426.63 (174.14 - 578.68) | 348.03 (53 - 565.03) | 358.975 (217.07 - 1636.62) | 302.015 (128.75 - 1194.23) | 332.9 (53.75 - 546.9) | 304.27 (216.34 - 608.39) |
| TNF⍺ | 21.6 (10.08 - 52.63) | 38.65 (7.71 - 123.62) | 17.21 (0 - 62.82) | 18.15 (0 - 78.24) | 16.68 (0 - 49.09) | 18.72 (0 - 66.88) |
| TNFβ | 0 (0 - 7.52) | 9.56 (0 - 40.52) | 0 (0 - 30.34) | 0 (0 - 25.03) | 0 (0 - 19.66) | 0 (0 - 8.09) |

**Table S2. Statistical Analysis.** Statistical analysis of analyte concentrations (pg/ml), with medians and ranges for each among healthy donors (HD), people with HIV (PWH) on combination antiretroviral therapy (cART), and viremic individuals.

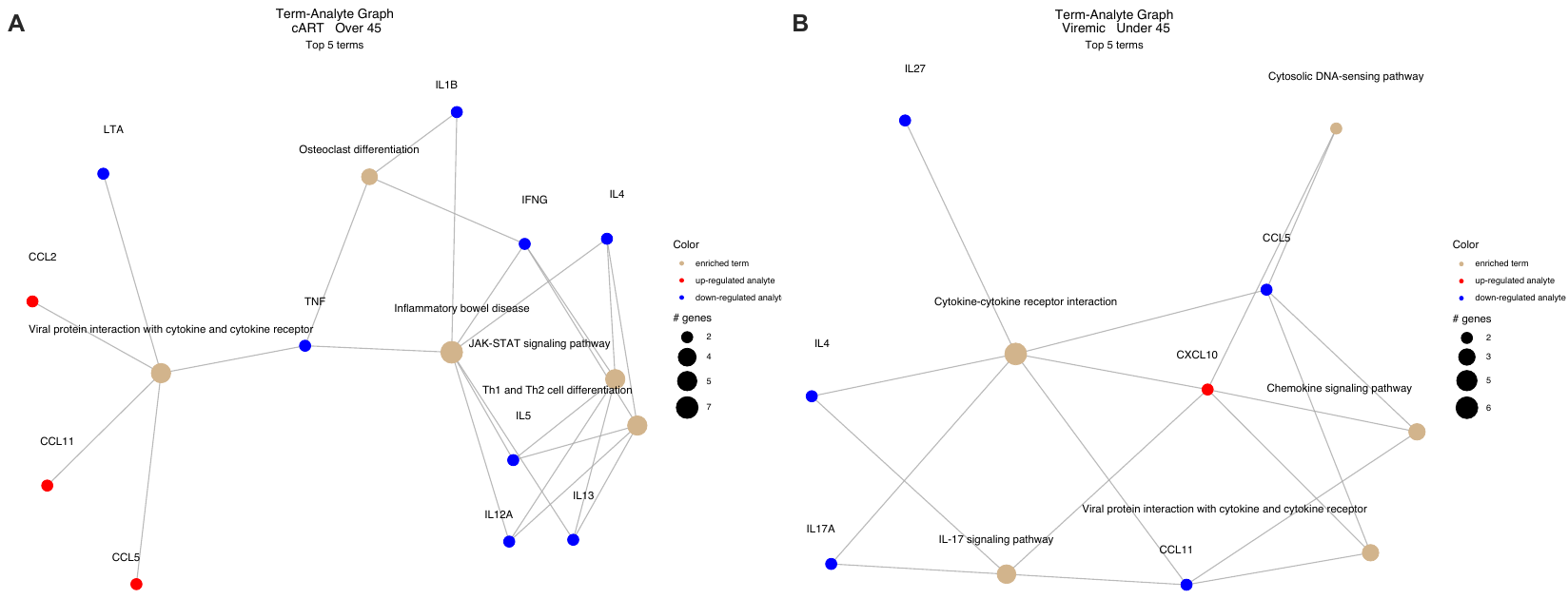

**Figure S3. Luminex Output.** Pathway analysis using term-analyte graphs for people with HIV (PWH) on combination antiviral therapy (cART) over 45 **(A)** and viremic individuals under 45 **(B)**. Data for pathway analysis was obtained by performing *t*-tests, and p-value cut-off of 0.2 was chosen. Fold changes for each group and age group were calculated compared to those of the healthy donor (HD) group and respective age group.
